# Backward collateral sensitivity can restore antibiotic susceptibility

**DOI:** 10.1101/2024.11.06.622341

**Authors:** Farhan R. Chowdhury, Brandon L. Findlay

## Abstract

The prevalence of antibiotic resistance continues to rise, rendering many valuable drugs ineffective. Antibiotic cycling regimens that incorporate collateral sensitivity (CS), the phenomenon where resistance to one antibiotic leads to hypersensitivity to another, are hypothesized to slow the evolution of antibiotic resistance. However, the repeatability of CS interactions and their ability to drive bacterial extinction and resensitizations remain unclear. In this study, we thoroughly investigate four drug pairs proposed for cycling regimens with experimental evolution. We find that reported pairwise CS interactions are not always robust, and even when they are, forward CS (where resistance to drug A leads to hypersensitivity to drug B) does not reliably reduce resistance or promote bacterial extinction. Instead, we find that if evolution of resistance to drug B in naive cells is associated with CS to drug A, a phenomenon we term backward CS, drug A-resistant cells can be rendered more sensitive to A again when resistance to B develops. We describe the mechanism of resistance disruption via backward CS in an aminoglycoside-β-lactam pair, where perturbation of the electron transport chain to inhibit aminoglycoside entry impairs β-lactam efflux. Overall, we highlight the importance of applying antibiotics in the correct order in cycling regimens and identify robust CS interactions that may be used to design treatment regimens less likely to lead to resistance evolution.

**TOC Graphic.**
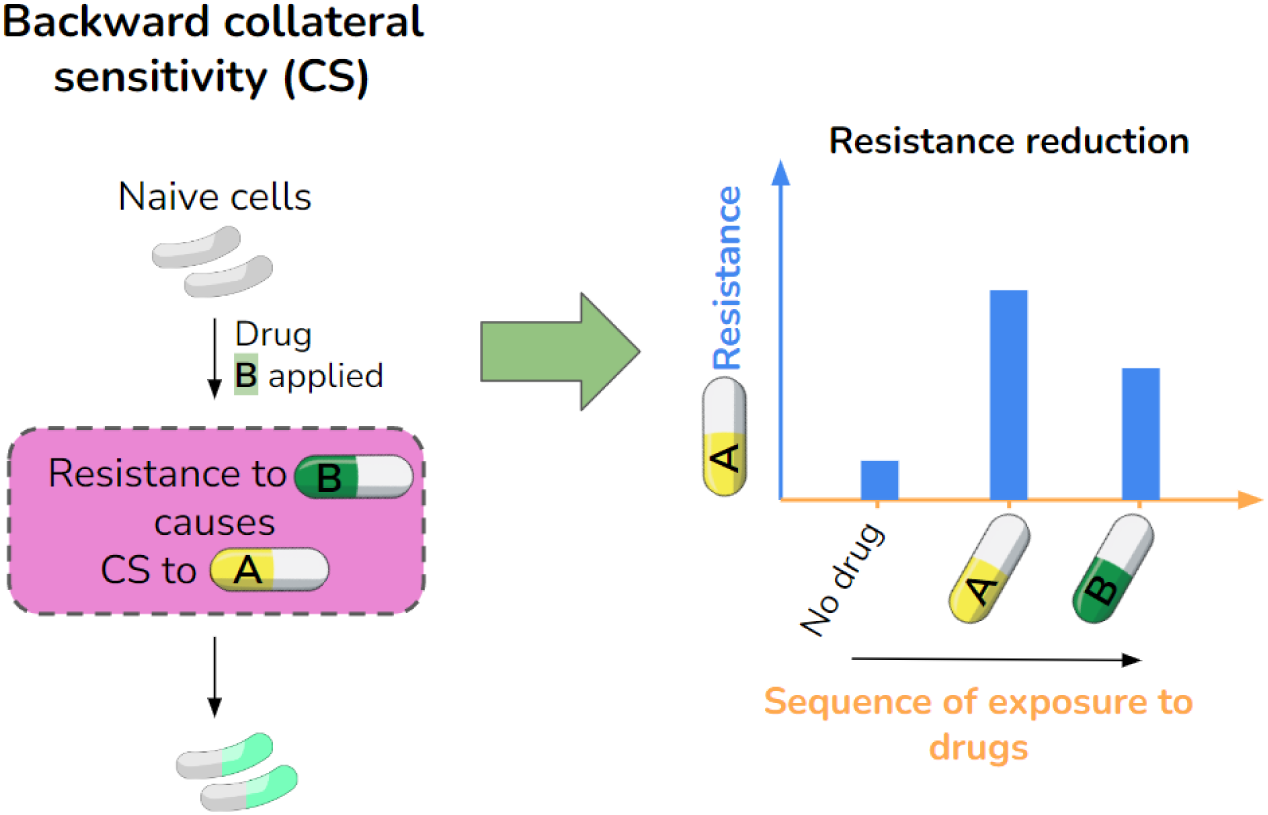

## Introduction

Antibiotic resistance is associated with 4.7 million deaths every year and is projected to claim 40 million lives by the year 2050 (1,2). As pathogens are adapting to antibiotics faster than new drugs can be developed, alternative strategies to curb resistance are necessary (3). One proposed strategy is to use existing drugs to design sequential or cyclic antibiotic treatment regimens with drugs applied one after the other at either defined time intervals or as resistance successively emerges (4–6). These sequential regimens are suggested to incorporate collateral sensitivity (CS) to enhance the efficacy of therapy by slowing or preventing the emergence of effective resistance (7–10). Collateral sensitivity is an evolutionary trade-off in bacteria where evolution of resistance to one antibiotic increases sensitivity to another. To date, work has focused on what we term *forward* CS, interactions where evolution of resistance to antibiotic A creates hypersensitivity to antibiotic B (11–13) (Figure 1). Subsequent exposure to antibiotic B is thought to then promote bacterial extinction and/or cause resensitization to antibiotic A (9). Unfortunately, to our knowledge such pairwise evolution work has only been completed with populations that all show CS to drug B, making it impossible to assess the role of CS in independently driving extinctions and resensitizations (9,14). Reports on the robustness of CS interactions are also conflicting, with some experimental evolution studies reporting that replicate populations exposed to one antibiotic frequently acquire CS to another (9,10,12,13), while others show weak reproducibility (15–17). Trade-offs must be evolutionarily repeatable to be useful in therapy (18), and given that antibiotic switching and compensatory evolution can affect resistance evolution independently of CS (14,19), it remains unclear if and to what degree CS disrupts evolution during antibiotic cycling.

**Figure 1:**
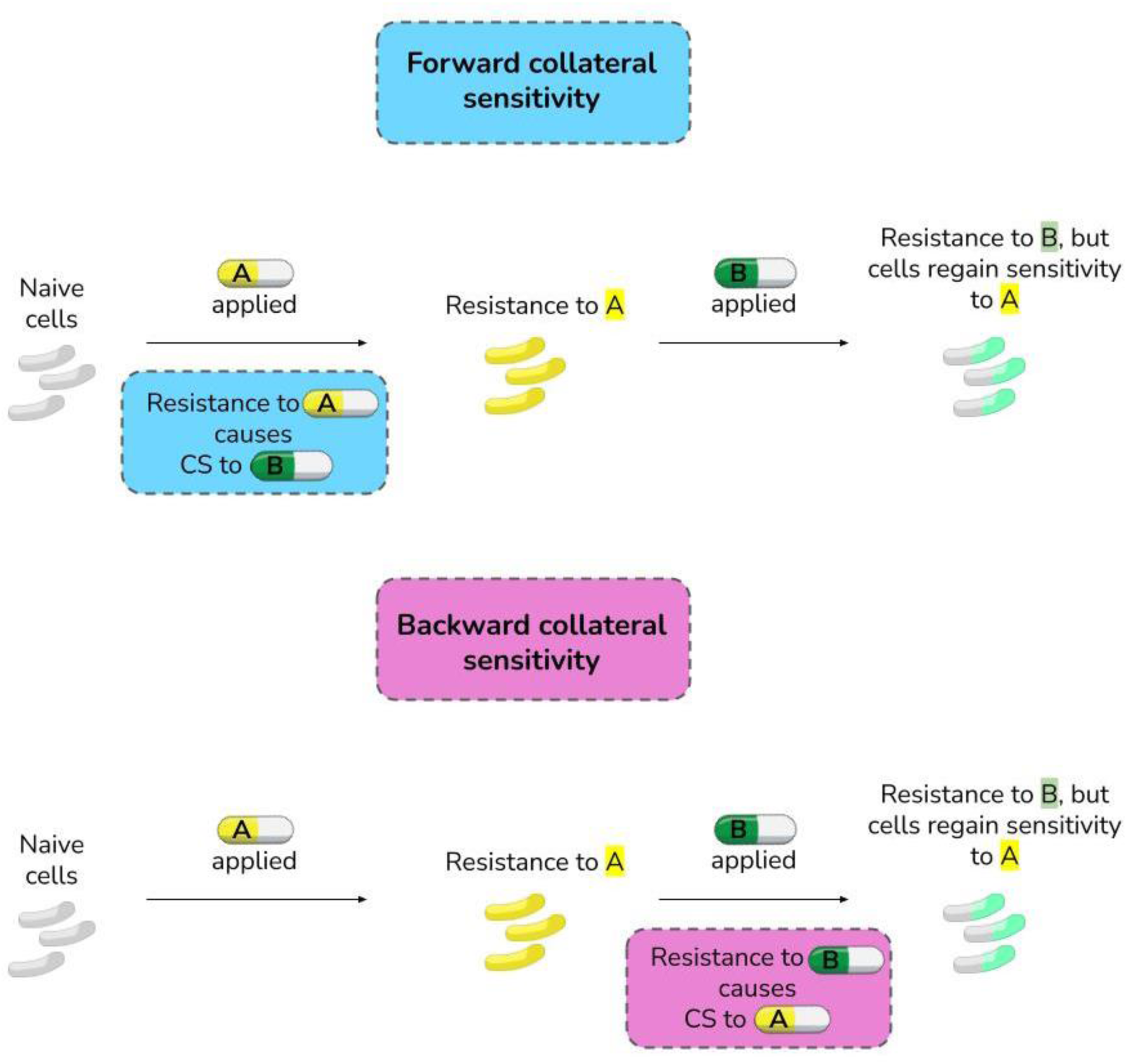
The concept of forward and backward collateral sensitivity.

In this work, we leverage the soft agar gradient evolution (SAGE) platform (6,20) to evolve 16 replicate populations of *Escherichia coli* K-12 substr. BW25113 against four drug pairs with reported CS interactions. We provide insight into the frequency of CS and cross resistance (CR) during adaptive laboratory evolution and, in combination with an evolution platform that selects for compensatory mutations (6), reveal the evolutionary stability of CS and its role in driving extinction and antibiotic resensitization. Among the 4 drug pairs tested, we find some that show robust CS interactions and some that show little to none. In the pairs that do show CS, forward CS does not significantly promote reduction in resistance to drug A or correlate with extinctions upon exposure to drug B in any pair tested. However, we find that if resistance to drug B in naive cells frequently produces CS to drug A, cells initially resistant to drug A are often rendered more susceptible to A when resistance to B emerges (Figure 1). We term this CS interaction *backward* CS (Figure 1), since CS to A evolves with resistance to B (CS direction: B to A) but antibiotics are applied in a sequence of A to B. We illustrate this by showing that in the aminoglycoside-β- lactam pair gentamicin (GEN) and piperacillin (PIP), GEN resistant populations exhibit widespread forward CS towards PIP, but subsequent exposure of cells with PIP CS to PIP does not lead to increased extinctions or significant GEN resistance reduction. However, when we expose PIP resistant bacteria to GEN (PIP – GEN pair; frequent CS in the opposite direction: backward CS), we see a 2-fold reduction in median PIP resistance. We use whole genome sequencing and efflux measurements to show that this reduction is driven by the weakening of PIP efflux in the GEN adapted strains. The effects of backward CS extend beyond the GEN – PIP pair, and we find polymyxin B (POL) – tigecycline (TIG) to be a pair that highly favors resensitization to POL. Our results show the importance of considering the direction of CS and drug switching in designing effective sequential therapies.

## Results

### A SAGE-based evolution platform to test for CS in pairwise drug sequences

We began with four drug pairs with previously reported CS between either the drugs or the drug classes: gentamicin (GEN) – piperacillin (PIP) (9), PIP – GEN (9,10), ciprofloxacin (CIP) – GEN (12), and polymyxin B (POL) – tigecycline (TIG) (10). First, we used SAGE to generate 16 independent replicates of *Escherichia coli* K-12 substr. BW25113 (WT) that were resistant to the first component of each drug pair at levels above clinical breakpoints (20) (Figure 2A). The sole exception was POL, where we generated 15 strains. High-level POL resistance was infrequent, and to generate 15 lineages required 88 starting replicates. Next, we passed these mutants through soft agar “flat plates” three times in series (Figure 2A). These plates contained the antibiotic from the prior challenge, at a concentration equal to half the minimum inhibitory concentration (MIC) of the antibiotic following SAGE. We included flat plates for three reasons: 1) general growth defects like slow growth rates, common after genomic adaptation to antibiotics (21) can appear as false CS during MIC plate readouts (10), 2) there were conflicting reports about the stability of CS (9,15), and 3) we wanted to study CS interactions that are not easily reverted via compensatory mutations. We previously showed that flat plates accelerate movement of chloramphenicol-resistant strains through soft agar, significantly improving growth rates in liquid media and allowing for resistance evolution to a subsequent antibiotic at near-wildtype frequencies (6). We find here that the effect is general, with similar effects on GEN resistant strains (Figure 2B). Replicates were then screened for resistance to the challenge antibiotic and for CS towards the second drug in the pair (Figure 2C, D). The majority of the strains cleared clinical breakpoints (22) for all the antibiotics tested (Figure 1C). WT MICs are listed in Table 1.

**Figure 2:**
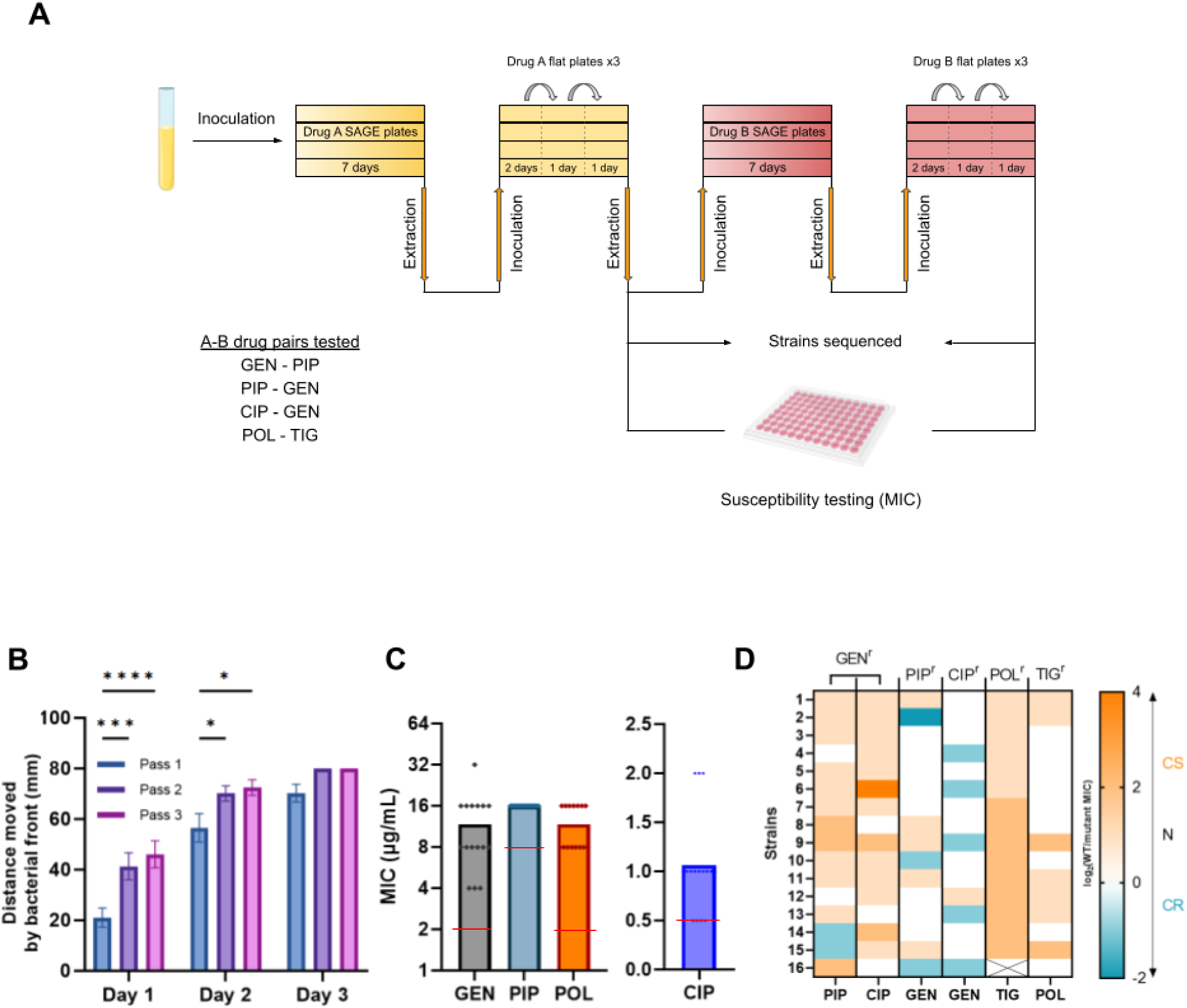
A SAGE-based evolution platform to study sequential antibiotic application. (A) Bacteria were inoculated in parallel into SAGE lanes containing antibiotic gradients, then incubated to generate resistant mutants. After 7 days, mutants were harvested and passed through three successive flat plates containing sub-inhibitory concentrations of the initial antibiotic. (B) Flat plates improve bacterial fitness in GEN resistant cells, as measured by distance swam (n= 16). (C) Following SAGE bacteria are resistant at or above clinical breakpoints. Red lines indicate the resistance breakpoints (n= 15 for POL, n= 16 for all other antibiotics). (D) Heatmap showing CS interactions between drugs. Bacteria are resistant to the drug labelled on the top of a column, and the label on the bottom shows CS readouts towards that drug. CS and CR are shown on a log2 scale. *p<0.05, ***p<0.001, ***p<0.0001, two-way ANOVA with Tukey’s multiple comparisons test.

**Table 1:**
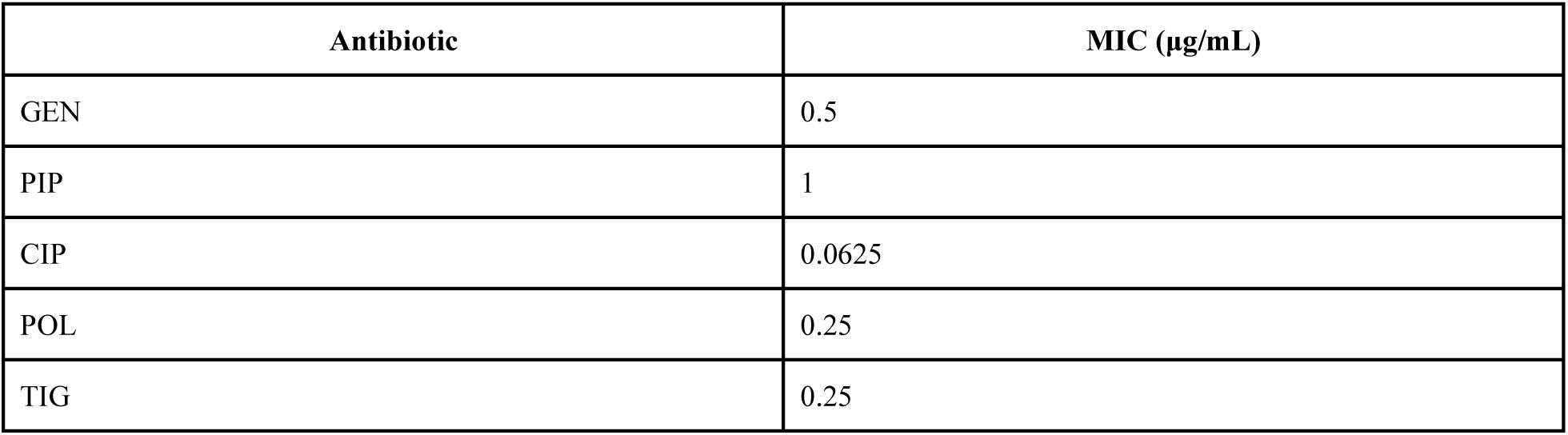
WT MICs.

| Antibiotic | MIC (µg/mL) |
| --- | --- |
| GEN | 0.5 |
| PIP | 1 |
| CIP | 0.0625 |
| POL | 0.25 |
| TIG | 0.25 |

PIP CS in GEN resistant replicates and TIG CS in POL resistant replicates occurred frequently, with little to no cross-resistance (CR) (Figure 2D). PIP resistant strains showed moderate GEN CS, while CIP resistant strains showed GEN CS in only 1/16 replicates. Our GEN – PIP CS results reinforce previous reports that aminoglycoside – β-lactam pairs exhibit reciprocal CS (9,10), but some reported CS interactions, such as between CIP – GEN (12), may either be infrequent or be mitigated via compensatory evolution.

### Forward CS does not promote extinctions, resistance drops or resensitizations in clonal populations

With a collection of strains with complete CS (POL – TOG), almost no CS (CIP – GEN), and a mix of both CS and CR (PIP – GEN and GEN – PIP) we then evaluated whether forward CS improves extinction rates and/or promotes resistance drops and resensitizations, by subjecting each resistant replicate to the second drug in its series (Figure 2A). We considered strains as resensitized to antibiotic A when both the following conditions were met: 1) resistance drops at least 4x from prior evolved MICs and 2) the MIC reduced to the clinical breakpoint or below. We set a strict definition for resensitization to accommodate for possible discrepancies due to random 2-fold MIC changes (23). Strains were considered extinct when cells could not be recovered after extraction from within 1.5 cm of the end of the SAGE plates.

We found that both the GEN – PIP and PIP – GEN pairs caused 8/16 of the replicates to go extinct (Figure 3A), even with significant differences in the prevalence of forward CS (Figure 2D). The extinctions occurred despite compensatory evolution in the flat plates, indicating a stable hurdle in adaptation to the second drug. The failures were not due to the antibiotic challenge alone, as generation of resistance to GEN or PIP in WT populations resulted in no extinctions (Figure 3A). The other two drug pairs did not show any extinctions, including the POL – TIG pair, which had ubiquitous forward CS (Figure 2D).

**Figure 3:**
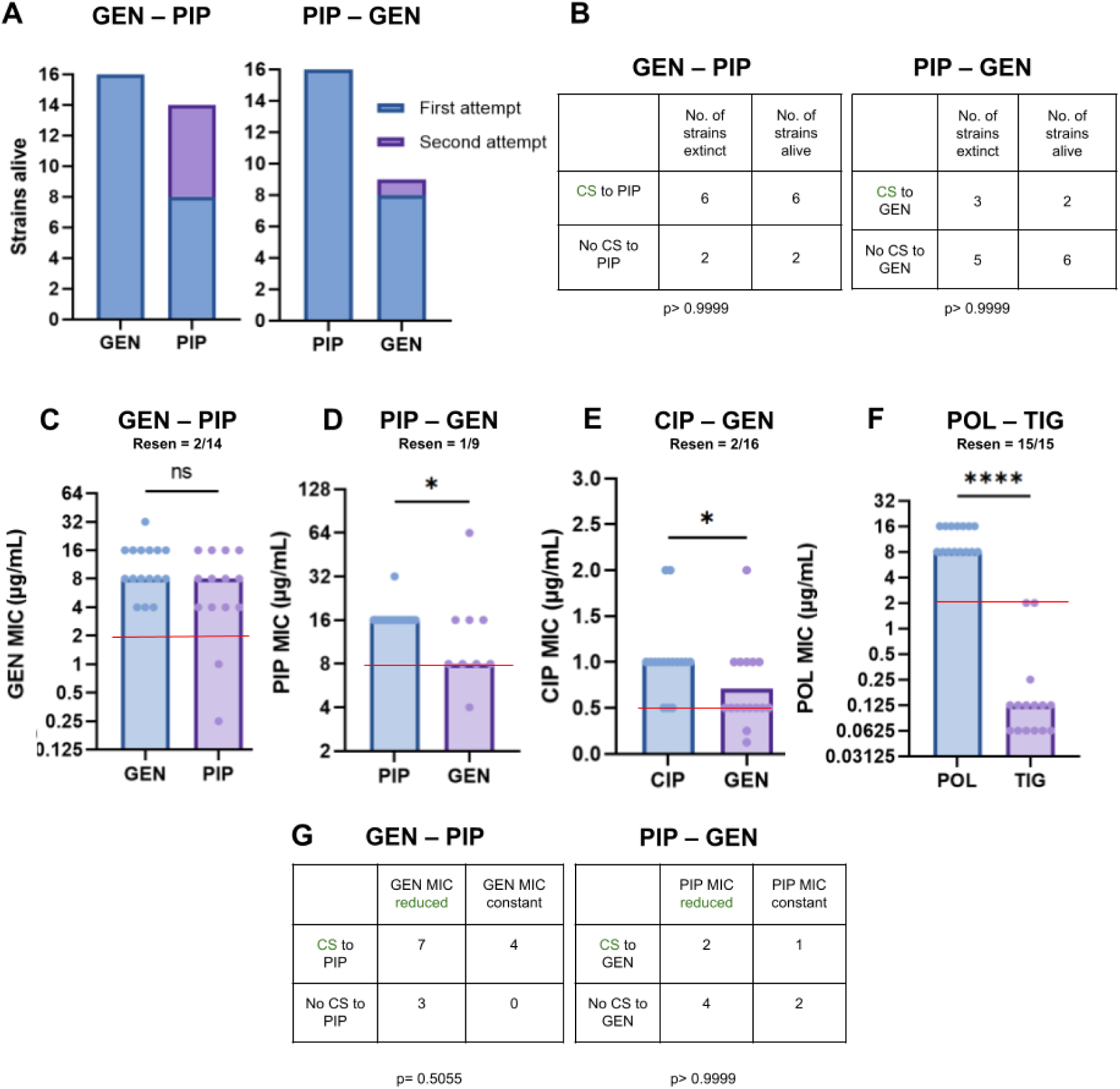
Extinctions, MIC reductions and resensitizations in drug pairs. (A) The GEN – PIP and PIP – GEN pairs cause frequent extinctions (n= 16 for both pairs). (B) Extinctions cannot be correlated with the incidence of CS (Fisher’s exact test). (C) The GEN – PIP pair does not cause a significant reduction in GEN MIC (n= 16 after evolution to GEN, n= 14 after evolution to PIP). (D) and (E) PIP – GEN and CIP – GEN cause a 2x reduction in median PIP and CIP resistance, respectively (n= 16 after evolution to PIP and CIP, n= 9 after evolution to GEN following PIP and n= 16 after evolution to GEN following CIP resistance). (F) The POL – TIG pair causes reliable resensitization and a large POL resistance drop (n= 15). Red lines indicate the resistance breakpoints. (F) and (G) Extinctions cannot be correlated with the incidence of CS (Fisher’s exact test). Resen = resensitizations. *p<0.05, ****p<0.0001, Mann-Whitney test.

To maintain the sample size for subsequent tests with the GEN – PIP and PIP – GEN pairs, extinct replicates were re-run using the same SAGE setup. This allowed recovery of 6/8 of the extinct replicates in the GEN – PIP pair, but only 1/8 in the PIP – GEN pair, the pair with lower incidence of forward CS (Figure 3A, 2D). We found no association between the replicates that went extinct and their CS status towards drug B in the GEN – PIP and PIP – GEN pairs (Figure 3B). This suggests that factors outside CS may drive extinctions in a sequential regimen.

To test the link between CS and reduced levels of resistance we first measured drug A resistance levels after exposure to drug B in the extant replicates. The GEN – PIP pair produced no significant drop in median resistance levels following PIP evolution, with 2/14 replicates resensitized to GEN (Figure 3C). The PIP – GEN and CIP – GEN pairs produced a 2-fold reduction in median drug A resistance, and 1/9 and 2/16 resensitizations respectively (Figure 3D, E). POL – TIG showed a remarkable 64x reduction in median POL resistance, achieving resensitizations in all 16 replicates (Figure 3F).

Next, we looked for associations between forward CS and drug A resistance drops. However, only 1/16 CIP resistant replicates showed GEN CS in the CIP – GEN pair, while all POL resistant replicates were resensitized to POL. Insufficient CS in the CIP – GEN pair, and the presence of only resensitized replicated in the POL – TIG pair made them unsuitable for this analysis. From the GEN – PIP and PIP – GEN pairs, we found no associations between the number of strains with reduced drug A resistance and forward CS (Figure 3G). This suggests forward CS may not play a significant role in resistance mitigation in a sequential regimen when clonal populations are involved. Overall, we found that the GEN – PIP and PIP – GEN pairs can cause reliable bacterial extinctions, with 3/4 drug pairs tested producing significant drug A resistance drops. However, extinctions and resistance drops were not associated with drug B CS, and resensitizations remained low.

### Backward CS may drive resistance drops

To explain the mechanism that drove the resistance drops we first looked into the PIP – GEN pair in which 5/9 replicates showed reduced PIP MIC after exposure to GEN (Figure 3C). GEN resistance is known to partially arise via mutations that weaken the proton motive force (PMF) (24) which may disrupt the efflux-driven PIP resistance (25). To test if PIP resistance is driven by efflux in our strains, we first sequenced three replicates after PIP exposure from the PIP – GEN pair (Figure 2A). Two of the three PIP-adapted strains showed mutations in genes known to affect efflux: *mprA* (26), *marR* (27), or *acrR* (28) (Figure 4A, Table 2A). All three strains also displayed CR to the antibiotics chloramphenicol (CHL) and tetracycline (TET) and the organic solvent hexanes. (Figure 4C, D, E). CHL, TET and hexanes are all known substrates of efflux pumps in *E. coli*, indicating increased efflux capacity in the PIP-adapted strains (29–31).

**Figure 4:**
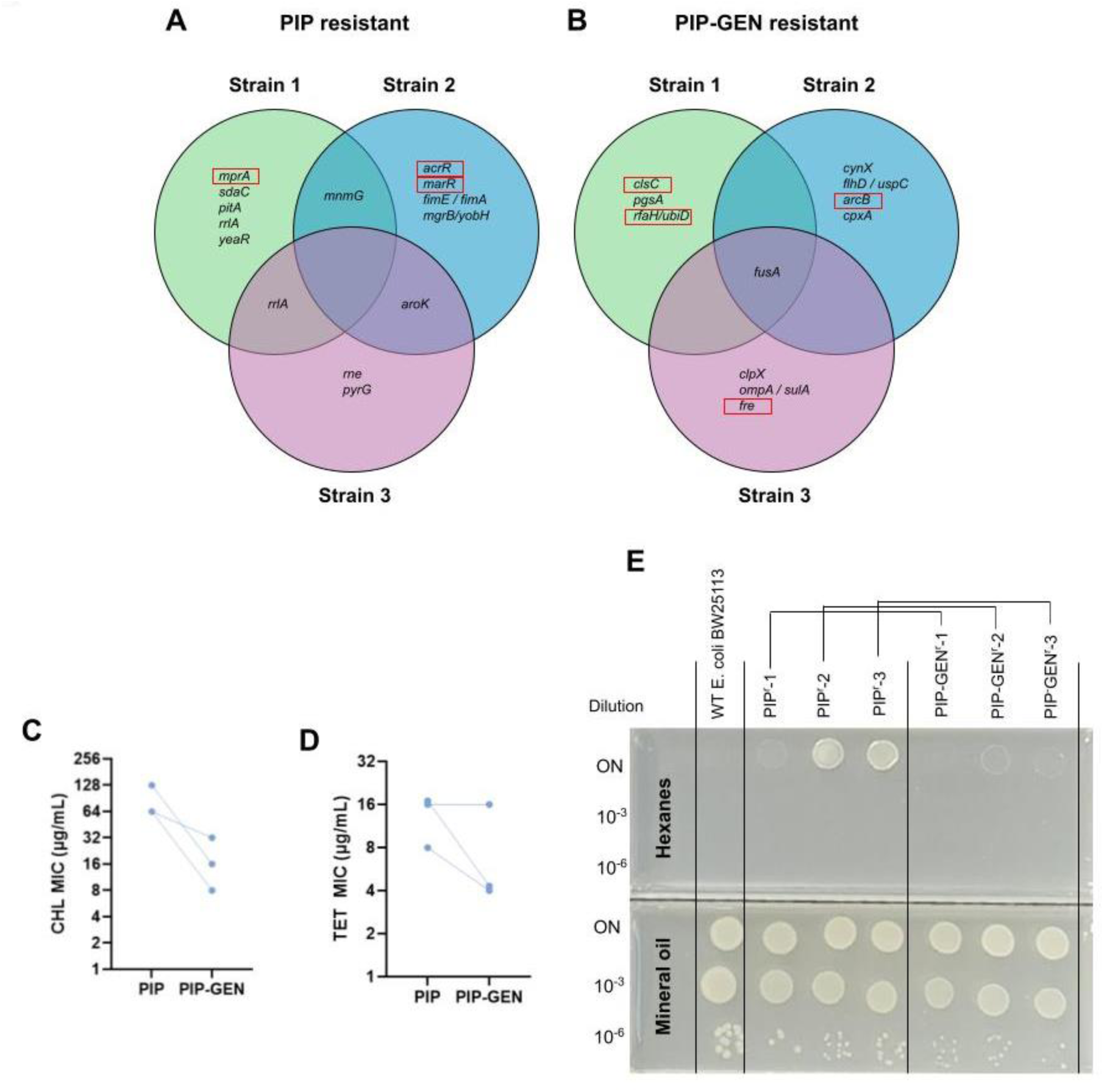
GEN resistance disrupts efflux-mediated PIP resistance. (A) and (B) Mutations identified in 3 strains. Genes boxed in red are involved in efflux activity or the electron transport chain (Table 2). (C) and (D) PIP-resistant strains are cross-resistant to CHL and TET, suggesting efflux upregulation in these strains. “PIP-GEN” strains that were sequentially adapted to PIP and GEN have on average increased CHL and TET susceptibility. (E) Hexanes tolerance test. All strains show good growth under mineral oil. The WT failed to grow under hexanes, while PIP resistant strains 2 and 3, and to a lesser extent, strain 1, showed growth on the undiluted spots. This ability is almost entirely lost upon GEN adaptation, suggesting efflux disruption. Representative picture from 3 independent experiments.

**Table 2:**
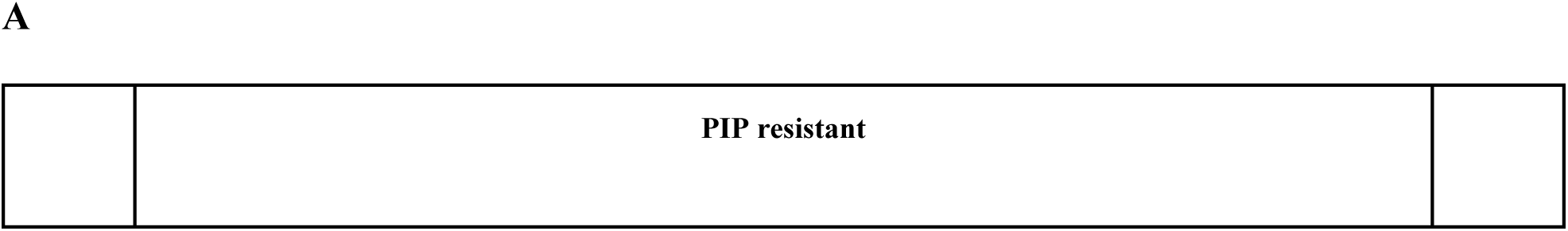
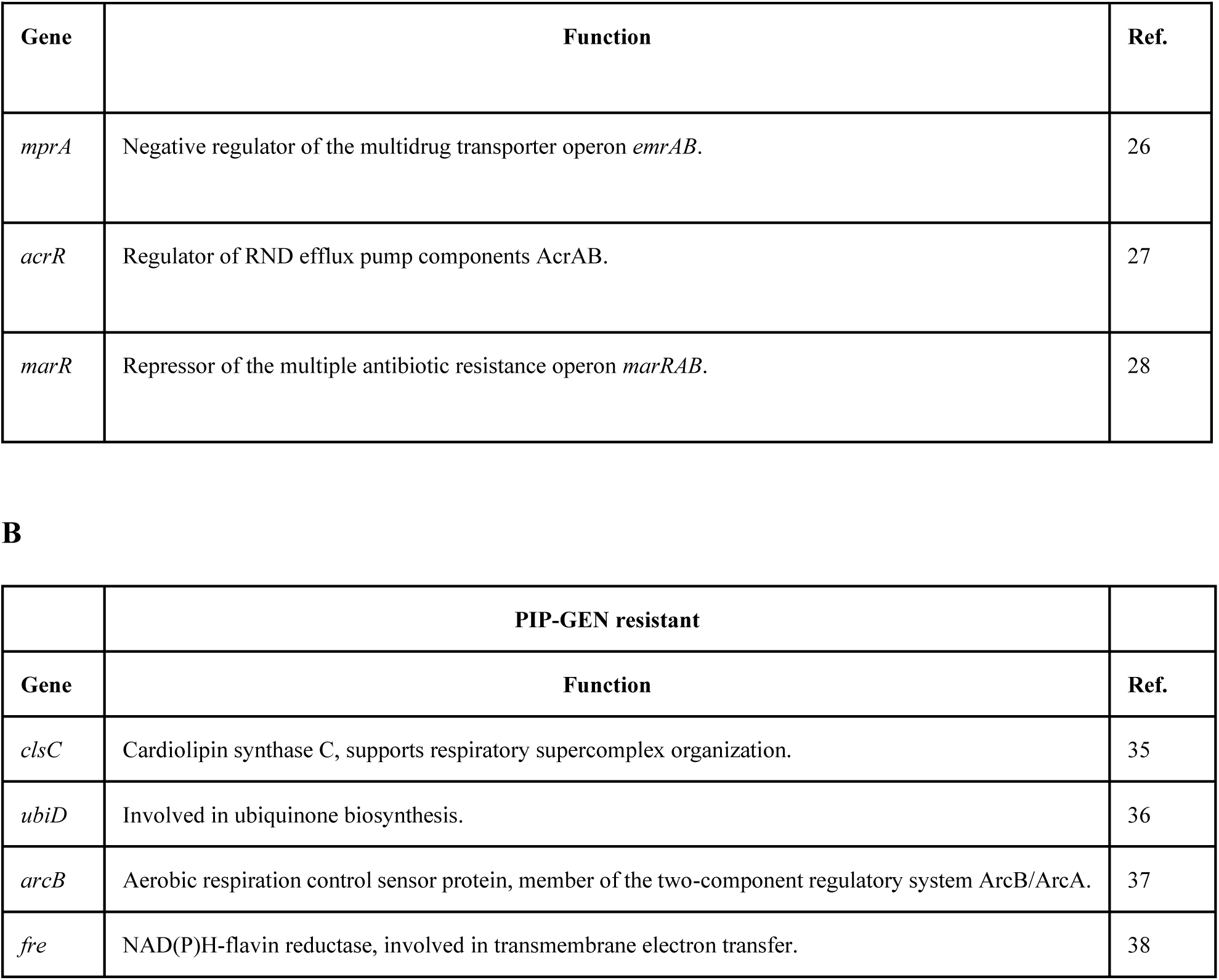
Mutations in the PIP and PIP-GEN adapted strains that affect efflux and the electron transport chain respectively.

| Gene | Function | Ref. |
| --- | --- | --- |
| *mprA* | Negative regulator of the multidrug transporter operon *emrAB*. | 26 |
| *acrR* | Regulator of RND efflux pump components AcrAB. | 27 |
| *marR* | Repressor of the multiple antibiotic resistance operon *marRAB*. | 28 |

**B**
|  | PIP-GEN resistant |  |
| --- | --- | --- |
| Gene | Function | Ref. |
| *clsC* | Cardiolipin synthase C, supports respiratory supercomplex organization. | 35 |
| *ubiD* | Involved in ubiquinone biosynthesis. | 36 |
| *arcB* | Aerobic respiration control sensor protein, member of the two-component regulatory system ArcB/ArcA. | 37 |
| *fre* | NAD(P)H-flavin reductase, involved in transmembrane electron transfer. | 38 |

To elucidate the effects of GEN resistance, we then sequenced three strains after GEN adaptation (Figure 2A). All three acquired mutations in *fusA* (Figure 4B), which codes for the elongation factor G and is known to confer gentamicin resistance (32). Additionally, every strain acquired mutations in genes involved in the electron transport chain: *clsC* (33), *ubiD* (34), *arcB* (35), or *fre* (36) (Table 2B). CHL resistance, TET resistance, and solvent tolerance all dropped following GEN adaptation (Figure 4C, D, E), showing that these mutations negatively affect efflux. The backward CS towards PIP that frequently arises with GEN resistance (Figure 2D, first column) may hence stem from a reduction in efflux. The median drop in PIP resistance levels after exposure to GEN also corresponds to the magnitude of backward PIP CS exhibited by WT cells resistant to GEN (Figure 2D first column, 3D).

To test if backward CS can also explain the resistance drops in CIP – GEN and POL – TIG pairs, we separately evolved 16 replicates against GEN and TIG to check the presence of backward CS towards CIP and POL respectively. GEN resistance imposed CIP CS in 13/16 strains (2x CS in 10, 4x CS one and 8x in two) (Figure 2D). Efflux is important in CIP resistance (37), and the efflux weakening effects of GEN resistance (Figure 4 C-E) could also be imparting the CIP CS. Again, the increase in CIP sensitivity in the CIP – GEN pair was equal to the magnitude of backward CIP CS (Figure 3E, Figure 2D, second column). Based on these results, we suggest that backward CS may disrupt resistance in A – B drug pairs, increasing drug A sensitivity. The effect of TIG resistance was more complex.

### POL resensitization is multifactorial

TIG resistance caused 2x POL CS in 5/16 strains and ≥4x CS in 2/16 strains (our limit of detection was 0.0625 μg/mL; MIC assays with wells clear at 0.0625 μg/mL were recorded as ≥4x CS) (Figure 2D, sixth column). Reciprocal CS between POL and TIG has been reported in *E. coli* before (10), but the mechanism of CS remains unknown. The resensitizations in the POL – TIG pair were generally stronger than the backward CS we observed, and with half of the POL – TIG replicates lacking CS (Figure 2D), backwards CS alone could not explain the median 64x increase in POL susceptibility. As POL MICs were measured after TIG SAGE evolutions and the flat plates (step 9 in Figure 5), it is possible that the change in susceptibility was due to heteroresistance or compensatory mutations (38). To identify the experimental stage at which the POL resensitizations occurred, we first revived four randomly selected POL resistant strains and passaged them five times through antibiotic free soft agar plates (Figure 5A, Supplementary figure 1). We measured POL MICs after each passage (Figure 5, **2** to **6**), and found a maximum 2x reduction in POL MICs (Figure 5B). Next, we revived the same four lineages from frozen stocks, but this time from stages 7 - 10 (Figure 5A) and measured their POL MICs. 1 out of the 4 strains was resensitized to POL during or immediately after evolution against TIG, while the rest showed a 0-4x reduction at this stage (Figure 5B). POL susceptibility in the other three strains was restored over the first TIG flat plate, which caused a 2-128x reduction in POL resistance. Subsequent TIG flat plates had no effect (Figure 5B). This suggests that cells with both POL and TIG resistance take evolutionary paths that promote phenotypic resistance reversion above what is achievable through the simple removal of POL selection pressure (11,19).

**Figure 5:**
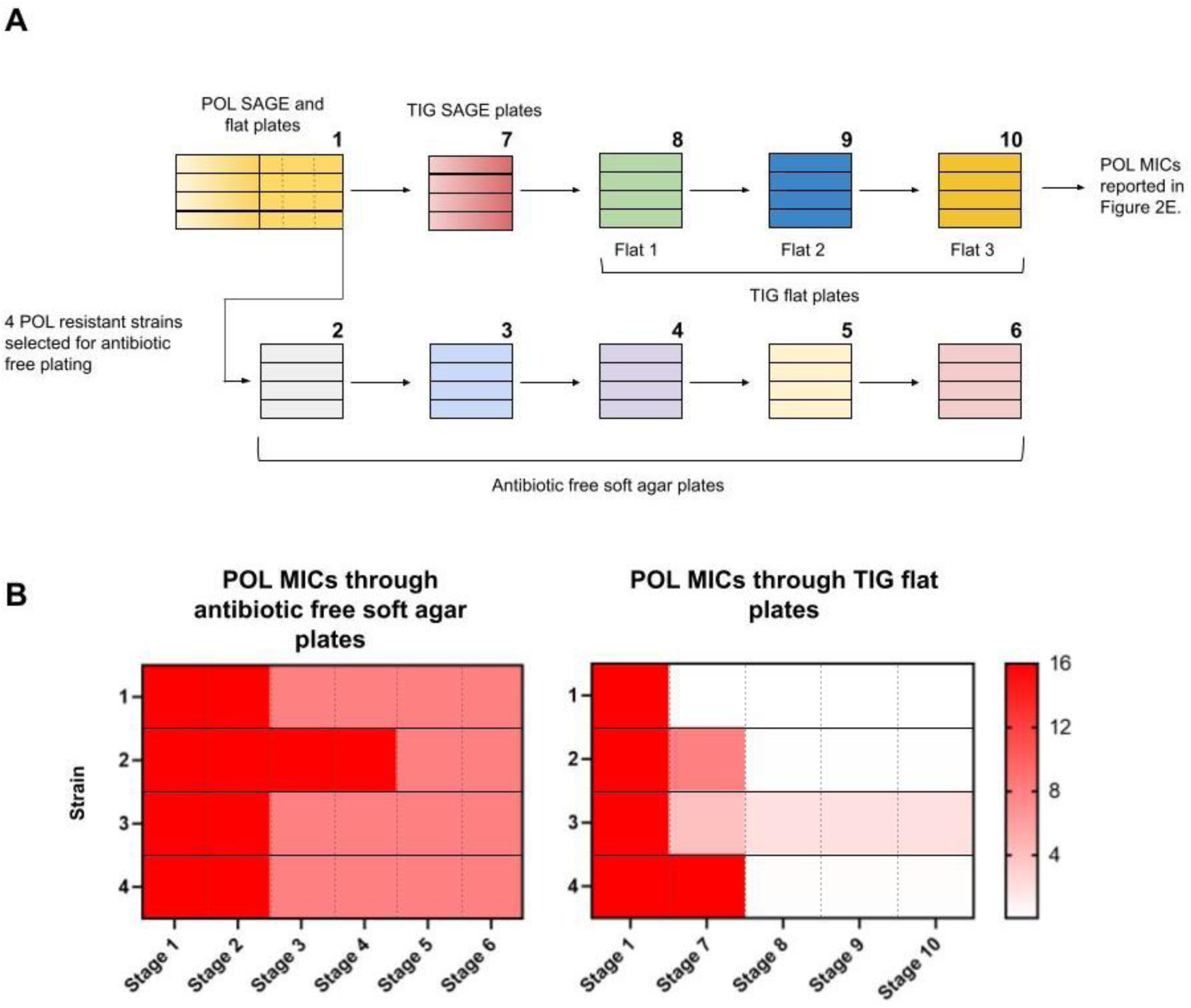
Resensitization of POL resistant strains. (A) Scheme showing the steps of POL and TIG sequential resistance evolution. Numbers represent the stages at which POL MICs were performed. (B) POL MICs at different stages of evolution.

## Discussion

In this study we investigated four drug pairs with previously reported CS interactions, generating 16 independent replicates of *E. coli* that were sequentially adapted to each drug in the pair. We observed a wealth of CS, CR and neutral interactions, allowing us to delineate the effect of CS on subsequent adaptive evolution and the resulting overall antibiotic susceptibility.

Contrary to prior reports (9,12,13), we found no significant associations between forward CS and bacterial extinction or antibiotic resensitization. Evolution of resistance to GEN led to frequent PIP CS (Figure 2D) but cells with and without PIP CS were equally likely to have reduced GEN resistance following evolution to PIP (Figure 3G). While PIP exposure did render half of the lineages extinct (Figure 3A), extinction was not correlated with the presence or absence of CS (Figure 3B).

In contrast, backward CS appeared to promote reduction resistance levels. Both the PIP – GEN and CIP – GEN pairs had high rates of backward CS and reductions in drug A resistance following the evolution of resistance to drug B (Figure 2D, Figure 3D, E). Our genomic analyses and efflux activity measurements showed that increased PIP sensitivity in the PIP – GEN pair was driven by the disruption of efflux capacity (Figure 4, Table 2), and may have also played a role in CIP CS. Backward CS was present, but not identified, in a prior study of CS pairs with reciprocal CS interactions (9), and our work suggests that backward CS may be partly responsible for the resensitizations seen in that study.

Although the GEN – PIP pair did not produce a statistically significant drop in median GEN MICs, the pair did produce individual replicates with reduced GEN resistance (Figure 3C, G). Median resistance levels were unaltered due to the presence of replicates with increased resistance (Mann-whitney test, p= 0.1515). Importantly, none of the drops in GEN resistance could be associated with PIP CS (Figure 3G). We also saw some backward CS in this pair (Figure 2D, third column), but the mechanism of aminoglycoside CS in β-lactam resistant bacteria is unknown. We sequenced a number of PIP resistant strains exhibiting a range of CS to GEN and observed mutations in genes involved in efflux and metabolism that could not be connected to GEN CS (data not shown).

All SAGE evolutions were conducted over a fixed period, with mutants extracted after seven days of incubation from within 1.5 cm of the end of the plates. This design allowed us to set a fixed benchmark to report adaptation rates and extinctions, but does not allow us to comment on if the speed at which resistance evolved was affected by the presence of CS. Future studies that track movement in SAGE plates could potentially answer this question.

The POL – TIG pair showed a 100% rate of POL resensitization, producing an impressive 64x drop in median POL resistance levels. While backward CS was also observed in this pair (Figure 2D, first reported by Imamovic *et al.* as reciprocal CS (10)), the magnitude of POL resistance reductions exceeded what would be expected from CS alone. Our results indicated that multiple mechanisms contributed to POL resensitization: removal of POL selection, adaptation to TIG, and probably most importantly, compensatory evolution in the POL and TIG resistant populations. This resensitization may be especially relevant for chronic diseases in which TIG and either POL or the polymyxin B analogue colistin see current use (39), such as cystic fibrosis (40).

Taken together, we suggest that backward, but not forward, CS may play an important role in reducing resistance levels in drug pairs. The weakening of efflux mechanisms upon exposure to a second antibiotic, as seen in the PIP – GEN, and possibly the CIP – GEN pair, may provide an approach for developing more effective sequential drug therapies aimed at reducing resistance. We also provide support for the idea that an aminoglycoside – β-lactam pair can frequently promote bacterial extinction. Our results highlight the importance of considering the directionality of CS interactions when designing sequential drug therapies and the need for thorough, large-scale screening for CS to build pairs more resilient against bacterial evolution.

## Materials and Methods

### Bacterial strain and growth conditions

E. coli K-12 substr. BW25113 and all subsequent resistant mutants were grown in Mueller Hinton (MH) media at 37 °C. Growth media was supplemented with appropriate antibiotics when growing mutants or extracting mutants from SAGE plates.

### SAGE Evolutions

SAGE plates were set up to generate resistant mutants as described before. All SAGE plates were made with Muller Hinton (MH) media + 0.15% agar + 0.2% xanthan gum (XAM). Antibiotic concentrations are listed in the table below, and were determined from trial SAGE experiments to evolve strains with MICs above clinical breakpoints within seven days. All SAGE plates were incubated for a fixed duration of seven days. Mutants were extracted from within 1.5 cm of the end of the lanes by pipetting 20 uL of the gel into Muller Hinton (MH) broth supplemented with the challenge antibiotic at a concentration = 2x the WT MIC (Table 1). Extracts were taken from regions with clear signs of growth. If no growth was apparent, extracts were pipetted from a random site within 1.5 cm of the end of the lane. A replicate was considered extinct if no growth was visible after overnight incubation in the antibiotic-supplemented MH broth.

| Antibiotic | Concentration (µg/mL) |
| --- | --- |
| GEN | 5 |
| PIP | 40 |
| CIP | 1 |
| POL | 10 |
| TIG | 5 |

### MIC assays

MICs were measured using the microdilution method outlined by the CLSI (41). Dilutions of antibiotics were prepared in MH broth and inoculated with bacteria at a final concentration = 1/200 of 0.5 McFarland standardized inoculum in non-treated 96-well plates. Plates were then incubated overnight and the MIC was recorded as the lowest concentration of antibiotic that prevented visible bacterial growth.

### Flat plates

Flat plates were prepared as previously described (6). First, the MIC of the antibiotic that was in prior SAGE plates was determined for all strains that completed their SAGE plates. Next, we created flat lanes specific for each strain by pouring ∼12 mL of XAM supplemented with the antibiotic at a concentration = ½ the MIC of that strain in a lane of a four-well dish. This allowed maintenance of the SAGE-evolved resistance phenotype during compensatory evolution. XAM media was used for all flat plates. Plates were inoculated as described before (20). Each replicate passed three consecutive flat lanes (Figure 2A, Figure 5). The first flat plate was incubated for two days, and the second and the third for one day (Figure 2A). The 16 GEN resistant strains were used to determine the appropriate flat plate incubation times, and all three passes for these strains were incubated for three days instead (Figure 3B). Extractions were carried out as described in the “SAGE evolutions” section, but were not limited to the 1.5 cm region of the end of the lanes. Instead, cells were extracted from where the cells had moved the farthest.

### Whole genome sequencing and analysis

Genomes were extracted from strains revived from frozen stock using the Bio Basic genomic DNA kit (Cat. no.: BS624). Sequencing and variant calling was performed by Seqcenter (USA). Sequencing was performed on an Illumina NextSeq 2000, and demultiplexing, quality control, and adapter trimming was performed with bcl-convert (v3.9.3). Variant calling was carried out using Breseq under default settings (80). NCBI reference sequence CP009273.1 for E. coli K-12 substr. BW25113 was used for variant calling. Figures showing common mutations in the three strains in Figure 4A and B were made using the R package *ggvenn*. For the PIP-GEN resistant strains, mutations acquired during PIP adaptation were removed before analysis.

### Hexanes tolerance assay

The solvent tolerance test was performed using a protocol adapted from Ikehata *et al* (42). Overnight cultures for each strain were diluted 10^3^ and 10^6^ times in MH broth and 5 μL spotted on MH agar surface. Spots were allowed to air dry, then the surface was either covered with ∼3 mm of hexanes (ACS Grade, Caledon Laboratory Chemicals, SKU: 5500-1-40) or mineral oil. Plates were sealed with parafilm and left in the fume hood to incubate for five days at room temperature. Plates needed refilling with hexanes every day due to evaporation.

### Antibiotic free soft agar plates

Plates were prepared similarly to the flat plates described before but without antibiotics. Strains were inoculated on one end of these plates and were incubated for one day. Strains were then extracted, cultured in antibiotic free MH broth and inoculated in a second plate. This process was repeated to achieve a total of five passes (Supplementary Figure 1). Broth cultured extracts were also streaked on antibiotic free MH petri plates. Cells from petri plates were used to perform MIC tests.

## Acknowledgments

This study was funded by the Fonds de recherche du Québec – Santé (FRQS) (269182). FRC is supported by the Fonds de recherche du Québec – Santé (FRQS) (B2X). We thank L. Freeman for helpful discussions.

---

**Supplementary Figure 1:**
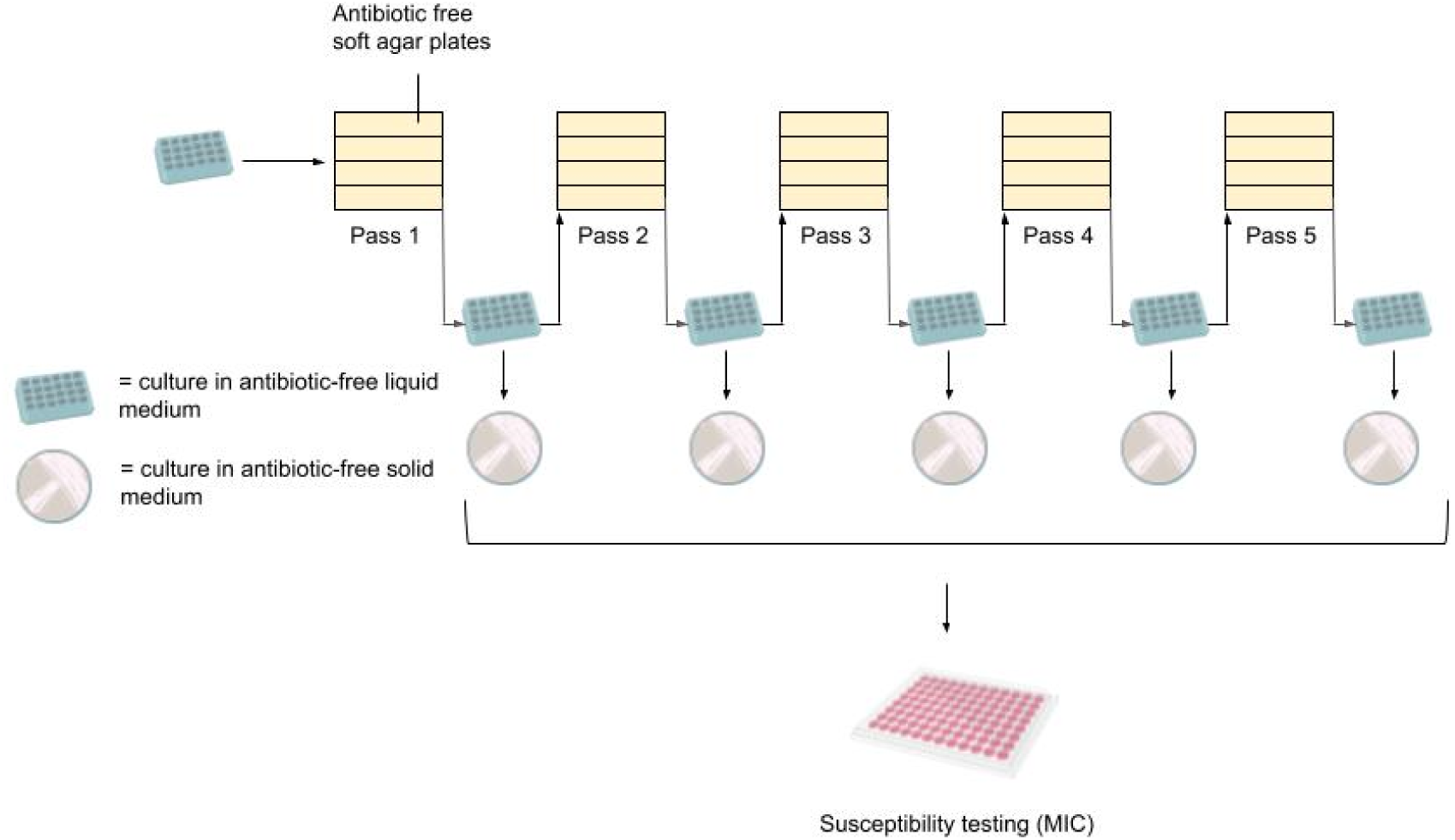
Schematic for antibiotic free soft agar passages. After the 5th pass strains have undergone 5x antibiotic free soft agar passages, 5x culturing in liquid without antibiotics, and 5x cultures on antibiotic-free solid media before being MIC tested.

## References

1. Murray CJL, Ikuta KS, Sharara F, Swetschinski L, Aguilar GR, Gray A, et al. Global burden of bacterial antimicrobial resistance in 2019: a systematic analysis. The Lancet. 2022 Feb 12;399(10325):629–55.

2. Naghavi M, Vollset SE, Ikuta KS, Swetschinski LR, Gray AP, Wool EE, et al. Global burden of bacterial antimicrobial resistance 1990–2021: a systematic analysis with forecasts to 2050. The Lancet. 2024 Sep 28;404(10459):1199–226.

3. Ventola CL. The antibiotic resistance crisis: part 1: causes and threats. P T Peer-Rev J Formul Manag. 2015 Apr;40(4):277–83.

4. Melnikov SV, Stevens DL, Fu X, Kwok HS, Zhang JT, Shen Y, et al. Exploiting evolutionary trade-offs for posttreatment management of drug-resistant populations. Proc Natl Acad Sci U S A. 2020 Jul 28;117(30):17924–31.

5. Andersson DI, Hughes D. Antibiotic resistance and its cost: is it possible to reverse resistance? Nat Rev Microbiol. 2010 Apr;8(4):260–71.

6. Chowdhury FR, Findlay BL. Fitness Costs of Antibiotic Resistance Impede the Evolution of Resistance to Other Antibiotics. ACS Infect Dis. 2023 Oct 13;9(10):1834–45.

7. Baym M, Stone LK, Kishony R. Multidrug evolutionary strategies to reverse antibiotic resistance. Science. 2016 Jan 1;351(6268):aad3292.

8. Hall MD, Handley MD, Gottesman MM. Is resistance useless? Multidrug resistance and collateral sensitivity. Trends Pharmacol Sci. 2009 Oct 1;30(10):546–56.

9. Barbosa C, Römhild R, Rosenstiel P, Schulenburg H. Evolutionary stability of collateral sensitivity to antibiotics in the model pathogen Pseudomonas aeruginosa. Wittkopp PJ, Pal C, Csorgo B, MacLean C, Johnsen P, editors. eLife. 2019 Oct 29;8:e51481.

10. Imamovic L, Sommer MOA. Use of Collateral Sensitivity Networks to Design Drug Cycling Protocols That Avoid Resistance Development. Sci Transl Med. 2013 Sep 25;5(204):204ra132–204ra132.

11. Roemhild R, Andersson DI. Mechanisms and therapeutic potential of collateral sensitivity to antibiotics. PLOS Pathog. 2021 Jan 14;17(1):e1009172.

12. Podnecky NL, Fredheim EGA, Kloos J, Sørum V, Primicerio R, Roberts AP, et al. Conserved collateral antibiotic susceptibility networks in diverse clinical strains of Escherichia coli. Nat Commun. 2018 Sep 10;9(1):3673.

13. Hernando-Amado S, Sanz-García F, Martínez J. Rapid and robust evolution of collateral sensitivity in Pseudomonas aeruginosa antibiotic-resistant mutants. 2020; Available from: https://www.semanticscholar.org/paper/0ce1a8541376ba7ed93464302e3414f548dc4290

14. Brepoels P, Appermans K, Pérez CA, Lories B, Marchal K, Steenackers H. Antibiotic cycling affects resistance evolution independently of collateral sensitivity. Mol Biol Evol. 2022 Dec 8;msac257.

15. Sørum V, Øynes EL, Møller AS, Harms K, Samuelsen Ø, Podnecky NL, et al. Evolutionary Instability of Collateral Susceptibility Networks in Ciprofloxacin-Resistant Clinical Escherichia coli Strains. mBio. 2022 Jul 7;13(4):e00441–22.

16. Nichol D, Rutter J, Bryant C, Hujer AM, Lek S, Adams MD, et al. Antibiotic collateral sensitivity is contingent on the repeatability of evolution. Nat Commun 2019 101. 2019 Jan 18;10(1):1–10.

17. Maltas J, Huynh A, Wood KB. Dynamic collateral sensitivity profiles highlight challenges and opportunities for optimizing antibiotic sequences. 2021; Available from: https://www.semanticscholar.org/paper/263c78d68029399947c9d431485b6e71dffd74af

18. Allen RC, Pfrunder-Cardozo KR, Hall AR. Collateral Sensitivity Interactions between Antibiotics Depend on Local Abiotic Conditions. mSystems. 2021 Nov 30;6(6):e01055–21.

19. Dunai A, Spohn R, Farkas Z, Lázár V, Györkei Á, Apjok G, et al. Rapid decline of bacterial drug-resistance in an antibiotic-free environment through phenotypic reversion. Landry CR, Wittkopp PJ, editors. eLife. 2019 Aug 16;8:e47088.

20. Ghaddar N, Hashemidahaj M, Findlay BL. Access to high-impact mutations constrains the evolution of antibiotic resistance in soft agar. Sci Rep 2018 81. 2018 Nov 19;8(1):1–10.

21. Melnyk AH, Wong A, Kassen R. The fitness costs of antibiotic resistance mutations. Evol Appl. 2015;8(3):273–83.

22. eucast: Clinical breakpoints and dosing of antibiotics [Internet]. [cited 2022 Oct 10]. Available from: https://www.eucast.org/clinical_breakpoints

23. Culp EJ, Waglechner N, Wang W, Fiebig-Comyn AA, Hsu YP, Koteva K, et al. Evolution-guided discovery of antibiotics that inhibit peptidoglycan remodelling. Nature. 2020 Feb;578(7796):582–7.

24. Lázár V, Singh GP, Spohn R, Nagy I, Horváth B, Hrtyan M, et al. Bacterial evolution of antibiotic hypersensitivity. Mol Syst Biol. 2013 Jan 1;9(1):700.

25. Seoane AS, Levy SB. Characterization of MarR, the repressor of the multiple antibiotic resistance (mar) operon in Escherichia coli. J Bacteriol. 1995;177(12):3414–9.

26. O L, K L, A M. EmrR is a negative regulator of the Escherichia coli multidrug resistance pump EmrAB. J Bacteriol [Internet]. 1995 May [cited 2024 Oct 14];177(9). Available from: https://pubmed.ncbi.nlm.nih.gov/7730261/

27. Beggs GA, Brennan RG, Arshad M. MarR family proteins are important regulators of clinically relevant antibiotic resistance. Protein Sci Publ Protein Soc. 2020 Mar;29(3):647–53.

28. Adler M, Anjum M, Andersson DI, Sandegren L. Combinations of mutations in envZ, ftsI, mrdA, acrB and acrR can cause high-level carbapenem resistance in Escherichia coli. J Antimicrob Chemother. 2016 May;71(5):1188–98.

29. Brunelle BW, Bearson BL, Bearson SMD. Chloramphenicol and tetracycline decrease motility and increase invasion and attachment gene expression in specific isolates of multidrug-resistant Salmonella enterica serovar Typhimurium. Front Microbiol. 2015 Jan 30;5:801.

30. Aono R, Tsukagoshi N, Yamamoto M. Involvement of Outer Membrane Protein TolC, a Possible Member of the mar-sox Regulon, in Maintenance and Improvement of Organic Solvent Tolerance of Escherichia coli K-12. J Bacteriol. 1998 Feb 15;180(4):938–44.

31. Pourahmad Jaktaji R, Zargampoor F. Expression of TolC and Organic Solvent Tolerance of Escherichia Coli Ciprofloxacin Resistant Mutants. Iran J Pharm Res IJPR. 2017;16(3):1185– 9.

32. Evgrafov MR de, Faza M, Asimakopoulos K, Sommer M. Systematic Investigation of Resistance Evolution to Common Antibiotics Reveals Conserved Collateral Responses across Common Human Pathogens. 2020; Available from: https://www.semanticscholar.org/paper/e7418f50efdf740c27e08def3219d824eecc2fea

33. Hryc CF, Mallampalli VKPS, Bovshik EI, Azinas S, Fan G, Serysheva II, et al. Structural insights into cardiolipin replacement by phosphatidylglycerol in a cardiolipin-lacking yeast respiratory supercomplex. Nat Commun. 2023 May 15;14(1):2783.

34. Gulmezian M, Hyman KR, Marbois BN, Clarke CF, Javor GT. The role of UbiX in *Escherichia coli* coenzyme Q biosynthesis. Arch Biochem Biophys. 2007 Nov 15;467(2):144–53.

35. Georgellis D, Kwon O, Lin EC. Quinones as the redox signal for the arc two-component system of bacteria. Science. 2001 Jun 22;292(5525):2314–6.

36. Schröder I, Johnson E, de Vries S. Microbial ferric iron reductases. FEMS Microbiol Rev. 2003 Jun 1;27(2–3):427–47.

37. Shariati A, Arshadi M, Khosrojerdi MA, Abedinzadeh M, Ganjalishahi M, Maleki A, et al. The resistance mechanisms of bacteria against ciprofloxacin and new approaches for enhancing the efficacy of this antibiotic. Front Public Health. 2022 Dec 21;10:1025633.

38. Andersson DI, Nicoloff H, Hjort K. Mechanisms and clinical relevance of bacterial heteroresistance. Nat Rev Microbiol. 2019 Aug;17(8):479–96.

39. Entenza JM, Moreillon P. Tigecycline in combination with other antimicrobials: a review of in vitro, animal and case report studies. Int J Antimicrob Agents. 2009 Jul;34(1):8.e1-9.

40. Parkins MD, Elborn JS. Newer antibacterial agents and their potential role in cystic fibrosis pulmonary exacerbation management. J Antimicrob Chemother. 2010 Sep 1;65(9):1853–61.

41. CLSI. M07: Dilution AST for Aerobically Grown Bacteria - CLSI [Internet]. 2018 [cited 2020 Jan 12]. Available from: https://clsi.org/standards/products/microbiology/documents/m07/

42. Ikehata Y, Doukyu N. Improving the organic solvent tolerance of *Escherichia coli* with vanillin, and the involvement of an AcrAB-TolC efflux pump in vanillin tolerance. J Biosci Bioeng. 2022 Apr 1;133(4):347–52.

